# Impact of Axon Model Complexity on Deep Brain Stimulation: A Comparative Analysis of MRG and Cohen Double-Cable Models

**DOI:** 10.64898/2026.08.23.746536

**Authors:** Mohadeseh Nadimi, Anne Rijpma, Ronald Bartels, Saman Vinke

## Abstract

Deep brain stimulation (DBS) modeling relies heavily on biophysical neuron models to estimate neural activation thresholds and predict stimulation spread. In this study, we systematically compared a widely adopted axon model, the McIntyre–Richardson–Grill (MRG) model (Model I), with a more detailed biophysical model, the Cohen model (Model II), to assess how structural and electrophysiological differences affect predicted DBS outcomes.

Electric field distributions generated by 2202 DBS lead were applied to the neuron models as extracellular input stimuli. Both models were simulated under biphasic pulse stimulation across varying axon–electrode distances, pulse widths, and stimulation frequencies.

Activation distances ranged from approximately 2 to 10 mm depending on stimulation parameters and contact location. At 2 mA, Model I achieved an activation distance of 6 mm, whereas Model II reached 10 mm, indicating greater excitability. Across matched fiber tracts, threshold differences ranged from −1.40 mA to 0.27 mA, with Model II requiring lower thresholds in 97.7% of cases. Both models showed a strong inverse relationship between pulse width and activation threshold. However, frequency responses differed: Model II exhibited increasing thresholds at higher frequencies, while Model I showed a slight decrease.

Machine learning regressors trained on distance, pulse width, and frequency achieved high predictive accuracy, with Gradient Boosting performing best. Model II demonstrated superior prediction metrics (R² = 0.986; RMSE = 0.045 mA; MAE = 0.034 mA) compared to Model I (R² = 0.977; RMSE = 0.089 mA; MAE = 0.068 mA).

Overall, both models reliably estimate DBS-induced activation, but structural differences significantly affect excitability and frequency-dependent behavior. With appropriate awareness of their respective strengths and limitations, either model can be used to derive activation distances for estimating electric field isolevels and the volume of tissue activated in patient-specific DBS simulations.

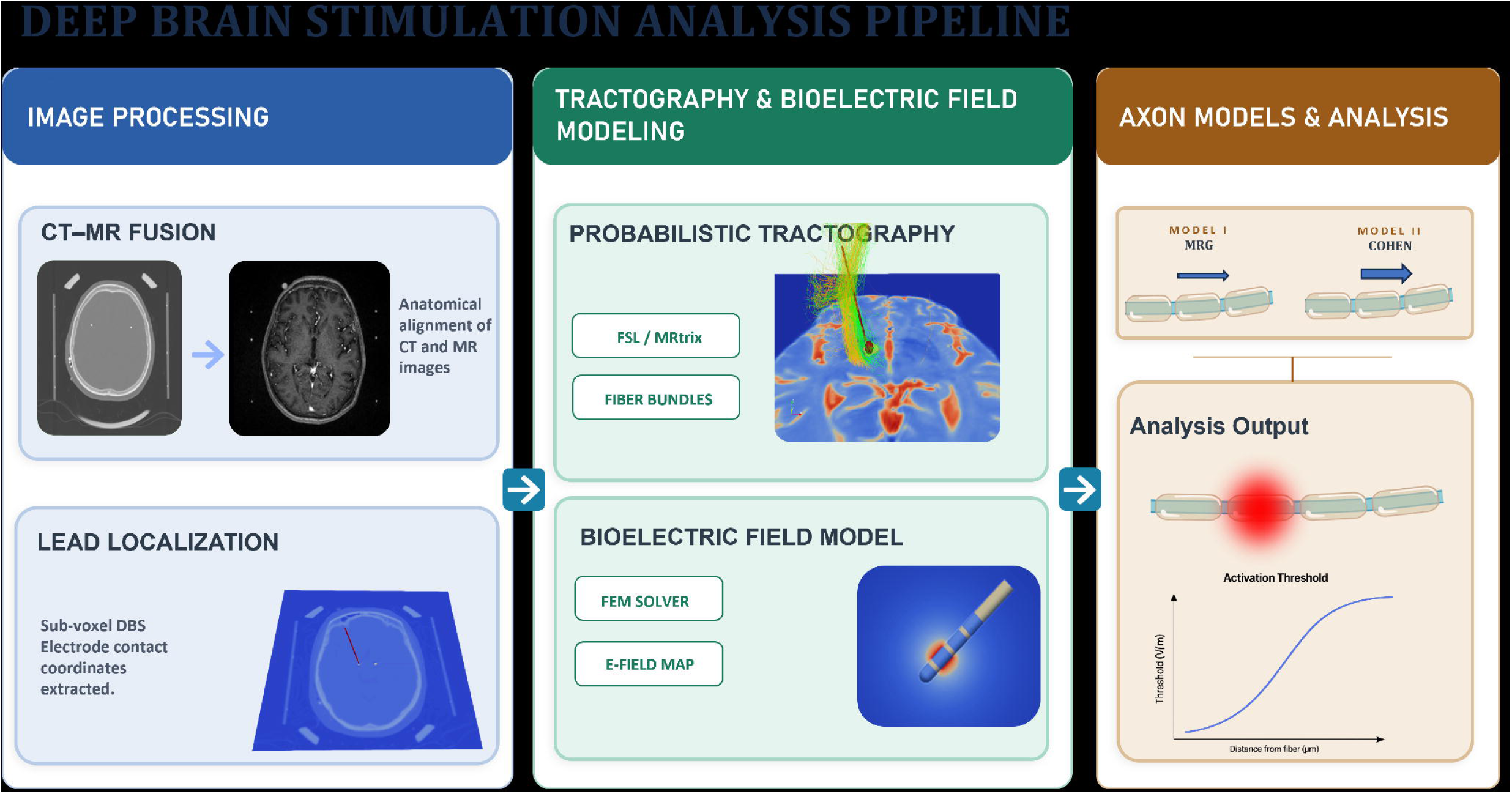

## 1. Introduction

Parkinson’s disease (PD) is a progressive neurodegenerative disorder characterized by disruptions in basal ganglia circuitry resulting from dopamine deficiency [1, 2]. In advanced stages, when pharmacological therapy becomes insufficient, deep brain stimulation (DBS) can provide effective and adjustable relief from motor symptoms [3, 4]. DBS delivers high-frequency (130 Hz) electrical pulses that modulate abnormal neural activity and help restore functional network dynamics [5, 6].

Recent advances in neuroimaging and electrode design have significantly improved the accuracy of DBS. Modern directional leads and current-steering systems allow for more selective targeting of brain structures, expanding therapeutic potential but also increasing the complexity of programming [7–10]. Determining optimal stimulation settings remains challenging due to inter-patient variability in electrode placement, stimulation parameters, and anatomical differences [11, 12].

To address these challenges, computational modeling has become a key tool for understanding and optimizing DBS effects. Patient-specific bioelectric field models enable estimation of the spatial distribution of stimulation-induced electric fields and prediction of neural activation patterns [13–15]. When coupled with axonal models, these simulations provide a biophysically informed estimate of the volume of tissue activated (VTA), a core metric linking stimulation parameters to clinical outcomes [16].

Among existing neuronal activation frameworks, the McIntyre-Richardson-Grill (MRG) model has been the standard for simulating myelinated axon responses to extracellular field [17–19]. While the MRG model employs a double cable geometry, it treats the submyelin pathway as largely passive and does not quantify periaxonal or paranodal resistivities. In contrast, Cohen et al. [20] introduced an experimentally constrained double cable model grounded in patch-clamp recordings, voltage sensitive dye imaging, and electron microscopy. Their work demonstrated that the periaxonal and paranodal spaces form a conductive nanocircuit that enables longitudinal current flow beneath the myelin and partially resistive coupling through paranodal junctions. This architecture reproduces submyelin potential gradients and explains temporal and amplitude saltation phenomena not captured by classical formulations. Incorporating this framework into DBS modeling could provide deeper insight into how stimulation waveforms interact with the microstructural properties of myelinated axons.

In this study, we explicitly implemented both the classical MRG axon model and the experimentally constrained Cohen double-cable model within the same DBS simulation framework. Using this controlled framework, we systematically quantified how the inclusion of a conductive periaxonal nanocircuit alters four key electrophysiological metrics: (i) membrane polarization along the axon during stimulation, (ii) activation threshold for action potential initiation, (iii) spatial recruitment patterns of axons surrounding the electrode, and (iv) sensitivity of activation to variations in stimulation amplitude, pulse width, and electrode-axon distance. Through this comparison, we aim to determine whether accounting for submyelin current flow provides a more physically realistic and predictive model of axonal activation during deep brain stimulation.

## 2. Methods

### 2.1 Patient Data and Image Processing

A single patient with Parkinson’s disease (female, 69 years old, 14 years since diagnosis) who underwent bilateral subthalamic nucleus (STN) DBS at Radboud University Medical Center was included in this study. Multimodal imaging data were acquired, including T1-weighted MPRAGE (FOV = 256 × 256 × 292 mm³, TR = 2300 ms, TE = 1.59 ms, flip angle = 8°, GRAPPA factor = 2) and T2-weighted sequence (FOV = 258 × 280 × 212 mm³, TR = 2500 ms, TE = 319 ms, and GRAPPA factor = 2), optimized for subcortical visualization and DBS target localization.

Structural images (T1-weighted MPRAGE and T2-weighted SPACE) were acquired on a 3T Siemens Skyra scanner (Siemens Healthineers, Erlangen, Germany) using an 18-channel UltraFlex head coil in combination with an 8-channel spine coil. These structural scans were obtained intraoperatively under general anesthesia to minimize motion artifacts.

Diffusion-weighted imaging (DWI) was obtained within five months prior to surgery using a single-shot spin-echo EPI sequence (b = 1000 s/mm², voxel size = 2 × 2 × 2 mm³, 76 slices, matrix = 120 × 120) with 64 non-collinear directions and both AP and PA phase-encoding polarities. DWI data were corrected for eddy currents and susceptibility-induced distortions. DWI acquisition was performed on a 3T Siemens Magnetom Prisma scanner (Siemens Healthineers, Erlangen, Germany) using a 20-channel Head/Neck coil.

A standardized image-processing workflow was implemented to construct a patient-specific DBS model using preoperative MRI and postoperative CT data. All analyses were performed in T1-weighted image space, with all other modalities co-registered accordingly.

Brain extraction was applied to both T1- and T2-weighted images using the ANTsPyNet deep-learning-based algorithm[21]. The extracted T1-weighted brain was segmented into cerebrospinal fluid (CSF), grey matter, and white matter using FMRIB’s Automated Segmentation Tool (FAST) [22]. The subthalamic nucleus (STN) was manually segmented by an expert based on T2-weighted MRI and the motor cortex was segmented using FreeSurfer [23].

Co-registrations were performed using ANTs, with the T1-weighted image serving as the reference space. The T2-weighted image was aligned to the T1 scan using the Symmetric Normalization (SyN) algorithm [24]. The resulting forward transform was applied to the manually segmented STN mask using nearest-neighbor interpolation.

The postoperative CT was cropped to remove inferior slices and neck regions, then rigidly registered to the T1-weighted MRI using FSL FLIRT [25] with a mutual-information cost function to handle cross-modality differences. Registration accuracy was confirmed using mutual information metrics and anatomical landmarks.

The registered CT image was used for electrode localization. The Pacer algorithm [26] automatically identified the DBS lead trajectory and position, while Diode v2 [27] automatically determined the rotational orientation of the directional lead, with the anterior direction defined as the 0° reference. Extracted coordinates were represented in RAS space; therefore, a sign-reversal of the x and y axes was applied to match the coordinate system required for VTK- based visualization.

Data were organized according to the Brain Imaging Data Structure (BIDS) standard [28], ensuring consistent file naming and metadata handling. Raw DICOM files were converted to NIfTI format using the dicom2nifti Python module.

### 2.2 Tractography

DWI data were pre-processed using MRtrix3 [29] for noise reduction and artifact correction, including denoising, Gibbs ringing removal, susceptibility distortion correction using opposing phase-encoded b₀ images, and correction for eddy currents, motion, and bias field inhomogeneities.

Targeted tractography was performed to reconstruct the hyperdirect pathway (HDP) rather than generating a whole-brain tractogram. Anatomically constrained tractography (ACT) was applied using a five-tissue-type (5TT) image to ensure biologically plausible streamline generation. Streamlines were seeded within Brodmann area 4 (primary motor cortex), defined by cortical parcellation, and constrained to pass through the manually segmented subthalamic nucleus (STN). The cerebral peduncle and the contralateral hemisphere were excluded to improve specificity.

To evaluate the influence of directional constraints on tract reconstruction, tractography was repeated with reversed inclusion and exclusion masks. This procedure served as a control to verify that the reconstructed hyperdirect pathway was not an artifact of the original masking strategy. By reversing the masks, we tested whether streamlines would still conform to the expected anatomical trajectory or if the reconstruction would produce implausible pathways, thereby confirming the specificity of the original approach.

Tracking parameters included a maximum curvature angle of 30° and the generation of 10,000 streamlines [30]. The resulting tractograms were refined to retain only streamlines within a physiologically relevant length range (10-100 mm) [31], ensuring that fibers traversed the STN and terminated within appropriate cortical and subcortical regions.

Regions of interest were transformed from T1/T2 anatomical spaces into diffusion space for anatomical correspondence, and tractography was performed in patient-specific diffusion space. Streamlines were linearly warped to T1-weighted space via a rigid-body registration (six degrees of freedom, normalized mutual information) using FLIRT. The transformation was converted to MRtrix format and applied to tractograms, which were subsequently exported to VTK for 3D visualization and analysis. This workflow ensured accurate spatial alignment between tractography and structural MRI.

### 2.3 Bioelectric Field Model

A volume conductor model was built from the imaging data through a structured, step-by-step modeling pipeline. First, a three-dimensional finite element mesh of the Boston Scientific DB- 2202 directional DBS lead and the surrounding brain tissue was generated using Gmsh [32]. The geometry was discretized with triangular surface and tetrahedral volume elements for accurate field computation in the lead and surrounding tissue. Curvature-based mesh refinement with a global characteristic length factor of 0.9 produced finer elements around the lead and coarser ones elsewhere, balancing accuracy and computational efficiency. The Frontal Delaunay and Delaunay algorithms were applied for 2D and 3D meshing, respectively, and automatic geometric coherence was enforced to ensure consistency in the final model. The brain volume was defined according to the dimensions of the T1-weighted image. The refinement process was iteratively performed until further mesh subdivision altered computed potentials by less than 1%, ensuring numerical convergence. The final mesh consisted of 132,532 triangular elements and 1,022,370 tetrahedral elements.

The electrode consisted of eight contacts, including two ring and six segmented directional contacts. Each contact had a height of 1.5 mm, separated by 0.5 mm of insulation, with an overall diameter of 1.3 mm. Physical groups defining individual surfaces and volumes were explicitly specified in the mesh to enable material assignment and boundary condition control within the FEM solver. The electrode position and orientation were determined from the postoperative CT scan, ensuring anatomically accurate alignment within the patient-specific head model. An encapsulation layer surrounding the electrode, with a thickness of 0.2 mm, was included to account for the low-conductivity fibrotic tissue that forms after implantation [18, 33]. The final mesh was exported from Gmsh and converted to Elmer FEM format using ElmerGrid [34].

Electrical properties were assigned to each material domain based on literature values for brain tissue, encapsulation, electrode, and insulation materials. The conductivity values were as follows: electrode = 1000 S/m, insulation = 0.001 S/m, brain tissue = 2 S/m and encapsulation tissue = 0.13 S/m [35].

The electric potential distribution was computed using the finite element method implemented in Elmer FEM under the quasi-static approximation (QSA), which neglects inductive and capacitive effects at deep brain stimulation (DBS) frequencies. The potential field in the conducting medium was obtained by solving the Laplace equation:

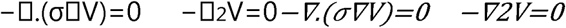

where V denotes the electric potential and σ the electrical conductivity. Conductivity was represented by a scalar value, assuming isotropic and frequency-independent behavior across all tissue domains.

For the patient-specific model, the clinically applied stimulation amplitude was imposed at the active electrode contact using a Neumann boundary condition, specifying the controlled current density to model current injection. A Dirichlet boundary condition was applied to the outer brain surface to represent the distant electrical ground. All remaining contacts were treated as floating (electrically inactive) boundaries, and zero-flux Neumann conditions were applied to all insulating surfaces to prevent current leakage and ensure physiologically realistic confinement of the electric field.

The steady-state conduction problem was solved using Elmer’s Static Current Solver, which computes the electric potential, electric field, and current density throughout the mesh. Numerical convergence and stability were ensured using an iterative BiCGStab solver with ILU preconditioning. Post-processing solvers were employed to calculate derived quantities such as electric field magnitude, current flux, and scalar outputs, which were exported in VTU and VTI formats for visualization and quantitative analysis. ParaView was used for visualization and quality inspection of the resulting mesh and potential fields.

### 2.4 Axonal models

To investigate how different axonal representations influence neural activation under DBS, both models (Model I (MRG) and Model II (Cohen)) were simulated using the NEURON environment (v8.2.2) [36] and Python (v3.10.12). Each model retained its original biophysical properties and geometry without modification, and identical stimulation protocols were applied to enable direct comparison of responses. Figure 1 provides a schematic comparison of the two formulations, highlighting their shared myelinated axonal architecture and the additional longitudinal conduction pathway in the periaxonal space incorporated in the Cohen model.

**Figure 1.**
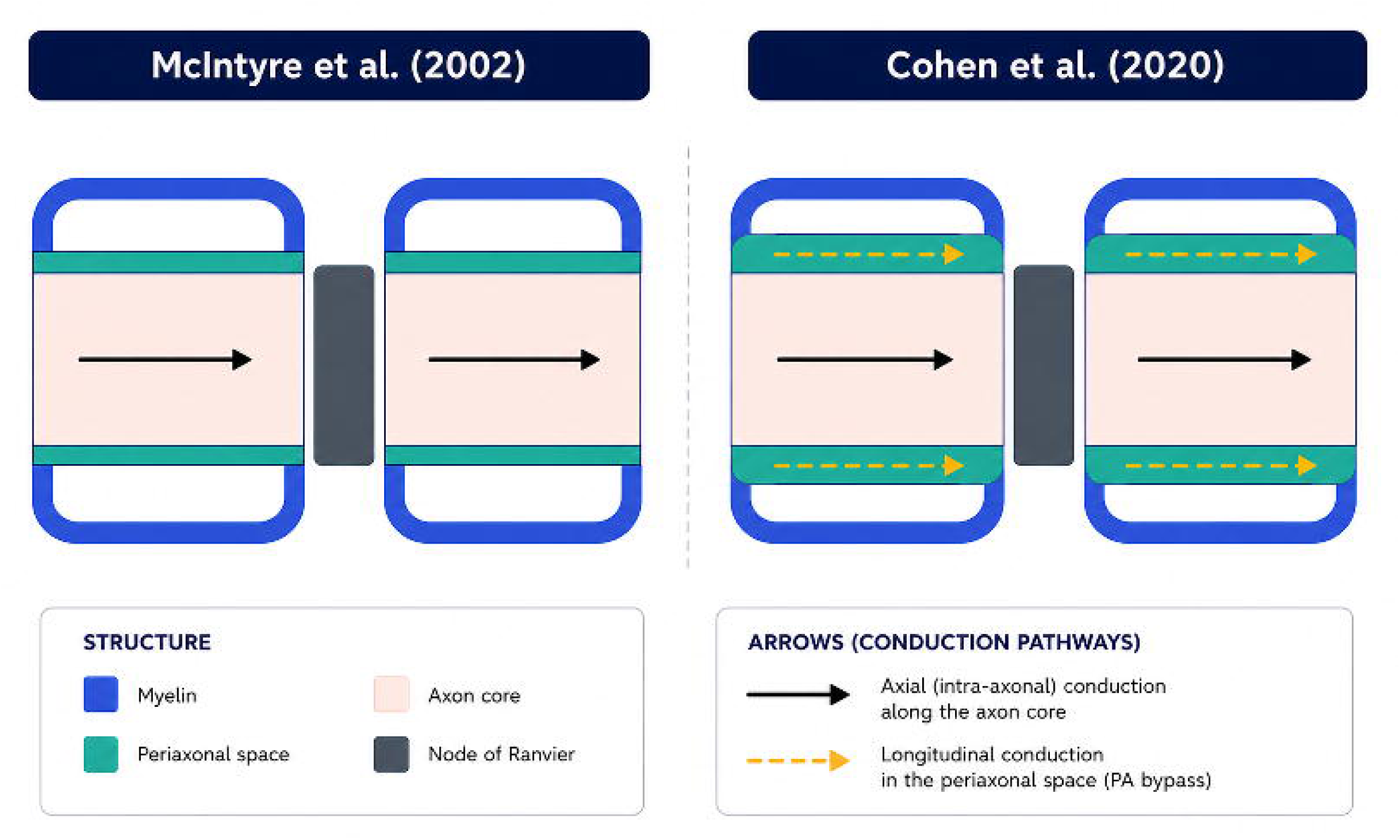
Schematic comparison of the double-cable axon models of McIntyre et al. (2002) and Cohen et al. (2020). Both models share the same double-cable architecture (myelin sheath, yellow; periaxonal space, green; axon, pink; node of Ranvier, grey) but differ in periaxonal (PA) space properties. In McIntyre et al. (2002) (left), the PA space is narrow and resistive; current flows through the axon core only (solid arrows), and PA discharge underlies the depolarising afterpotential. In Cohen et al. (2020) (right), the PA space is 12.3 nm wide and conductive (∼53.7 Ω·cm), acting as a longitudinal bypass pathway (dashed arrows) that enables the downstream node to depolarise ∼100 µs earlier temporal saltation.

#### 2.4.1 MRG model^1^

The MRG model with a 3 µm diameter was adopted to represent the DCF axon. This model was originally introduced in 2002 [17]and its geometric parameters were updated in 2016 [37]. In the MRG model, the axon contains 21 nodes of Ranvier with myelin attachment segments (MYSA), paranode main segments (FLUT) and internode segments (STIN) with total compartments of 221. Ionic conductances at each node are modelled using Hodgkin–Huxley-type mechanisms for fast sodium (Na⁺), persistent sodium, slow potassium (K⁺), and leakage currents [38].

#### 2.4.2 Cohen double-cable model^2^

We used a biophysically-detailed compartmental model of a layer 5 cortical pyramidal neuron [20]. The model consists of 95 myelinated axon sections with total 292 compartments (length range: 0.69 - 929.46μm, diameter range: 0.32- 4.70) extending from a multi-compartment soma and dendritic tree.

Each internode is represented by two coupled cables: one describing the axoplasm, and the other the periaxonal space bounded by the myelin sheath. Model parameters were derived by fitting experimental data from patch-clamp and voltage-sensitive dye recordings. These include periaxonal resistivity (53.7 Ω·cm), paranodal resistivity (550 Ω·cm).

#### 2.4.3 Simulation setup

One hundred axonal trajectories corresponding to the hyperdirect pathway (HDP) were extracted from the patient’s diffusion tractography data for simulation. A region of interest (ROI) surrounding the DBS lead, with dimensions of 20 × 19.5 × 20 mm was defined using the SaveGrid function at a resolution of 0.5 mm within the FEM solution domain (Section 2.3). The electric potential field was resampled onto this uniform fine grid to improve the spatial accuracy of extracellular potential mapping.

The stimulation waveform was modeled as an asymmetric charge-balanced, biphasic rectangular pulse train at the target frequency. Each pulse included a 0.06 ms cathodic phase at 1.0 mA followed by a 0.02 ms interphase gap and an anodic discharge phase with an amplitude 50 times smaller than the cathodic phase. The anodic component featured a prolonged exponential decay with a discharge duration of 3.7 ms and an RC time constant of 1 ms, ensuring net zero charge delivery per cycle.

The time-varying stimulation waveform was applied as a temporal scaling factor to the steady-state potential distribution obtained from the FEM model (section2.3), yielding the time-dependent extracellular potential experienced by each axon. This potential was then coupled to node compartments based on their spatial positions, via NEURON’s extracellular mechanism, simulating the effect of the surrounding electrical field on the axon’s membrane potential [39].

Action potential initiation was defined by threshold crossing −20 mv. An axon was considered activated if it generated five consecutive action potentials at the nodes of Ranvier originating from the stimulation site [40]. The first node was positioned adjacent to the stimulation site, consistent with the principle that axonal activation typically begins near the axon. Activation thresholds were determined using a binary search algorithm with a 1% tolerance, ensuring accurate convergence to the minimal stimulation amplitude required for activation. Each simulation used a time step of 1 µs and a total duration of 7.69 ms per stimulus pulse. All simulations were performed at 37 °C, and outputs were analyzed using custom Python scripts for post-processing and visualization.

### 2.5 Analysis and Comparison Metrics

To investigate how stimulation parameters influence action potential propagation, we conducted a series of simulations using both the Cohen and MRG axonal models. Clinically relevant ranges of pulse width and frequency were applied to assess model responses under DBS-like conditions.

The primary comparison metrics were as follows:

- Activation Distance (mm): Defined as the distance from the stimulation source (DBS lead) to the farthest axon that generates an action potential. This metric reflects the spatial extent of neural activation.
- Activation Threshold: The minimum stimulation amplitude required to elicit a propagating action potential, determined using a binary search algorithm (see Section 2.3.3).

Additionally, to explore how stimulation settings influence model behavior, the following parameters were varied:

- Pulse Width: 60 μs, 90 μs, and 120 μs
- Frequency: 80 Hz, 130 Hz, and 180 Hz

To predict activation thresholds from stimulation parameters, we implemented three regression models: Linear Regression, Random Forest, and Gradient Boosting. These models were selected to span a range of complexity and interpretability, from simple linear relationships to advanced ensemble methods capable of capturing nonlinear effects.

Linear Regression was used as a baseline model to quantify the linear relationship between predictors (distance, pulse width, frequency) and activation threshold.

Random Forest is an ensemble of 100 decision trees (n_estimators=100, random_state=42), which aggregates predictions to reduce overfitting and model nonlinear interactions. Feature importance scores were extracted to assess the relative influence of each parameter.

Gradient Boosting constructs an ensemble of 100 sequential trees (n_estimators=100, random_state=42), optimizing prediction accuracy by correcting errors of previous trees. This method is well-suited for tabular data and can model complex dependencies.

All input features were standardized using z-score normalization (StandardScaler) to ensure comparability and improve model convergence. Rows with missing values in any feature or the target variable were excluded from analysis. Models were trained on 80% of the data and tested on the remaining 20%. Performance was assessed using the coefficient of determination (R²), root mean squared error (RMSE), and mean absolute error (MAE) on the test set. To further validate generalizability, 5-fold cross-validation was performed on the training set, reporting the mean and standard deviation of R² scores. For tree-based models, feature importance was reported to quantify the contribution of each predictor.

Agreement between models for activation threshold was assessed using linear regression and Bland-Altman analysis. Paired threshold predictions from both models (n = 1051) were compared across all stimulation parameter combinations (pulse width: 0.06-0.12 ms; frequency: 80-300 Hz).

The coefficient of determination (R²), root mean squared error (RMSE), and mean absolute error (MAE) were calculated to quantify agreement. Pearson correlation was used to assess the strength of the linear relationship. Bland-Altman plots were generated to visualize systematic bias and limits of agreement.

## 3. Results

Figures 2 illustrate activation thresholds and activation distances for Models I and II across different contact points under biphasic pulse stimulation. For both models, activation distance ranged from approximately 2 mm to 10 mm. At 2 mA with a biphasic pulse, Model I achieved an activation distance of 6 mm, whereas Model II reached 10 mm. For matched tracts, the difference in activation threshold between models I and II ranged from −1.40 mA to 0.27 mA. Notably, model II demonstrated greater sensitivity, requiring a lower activation threshold for 97.7% of tracts.

**Figure 2.**
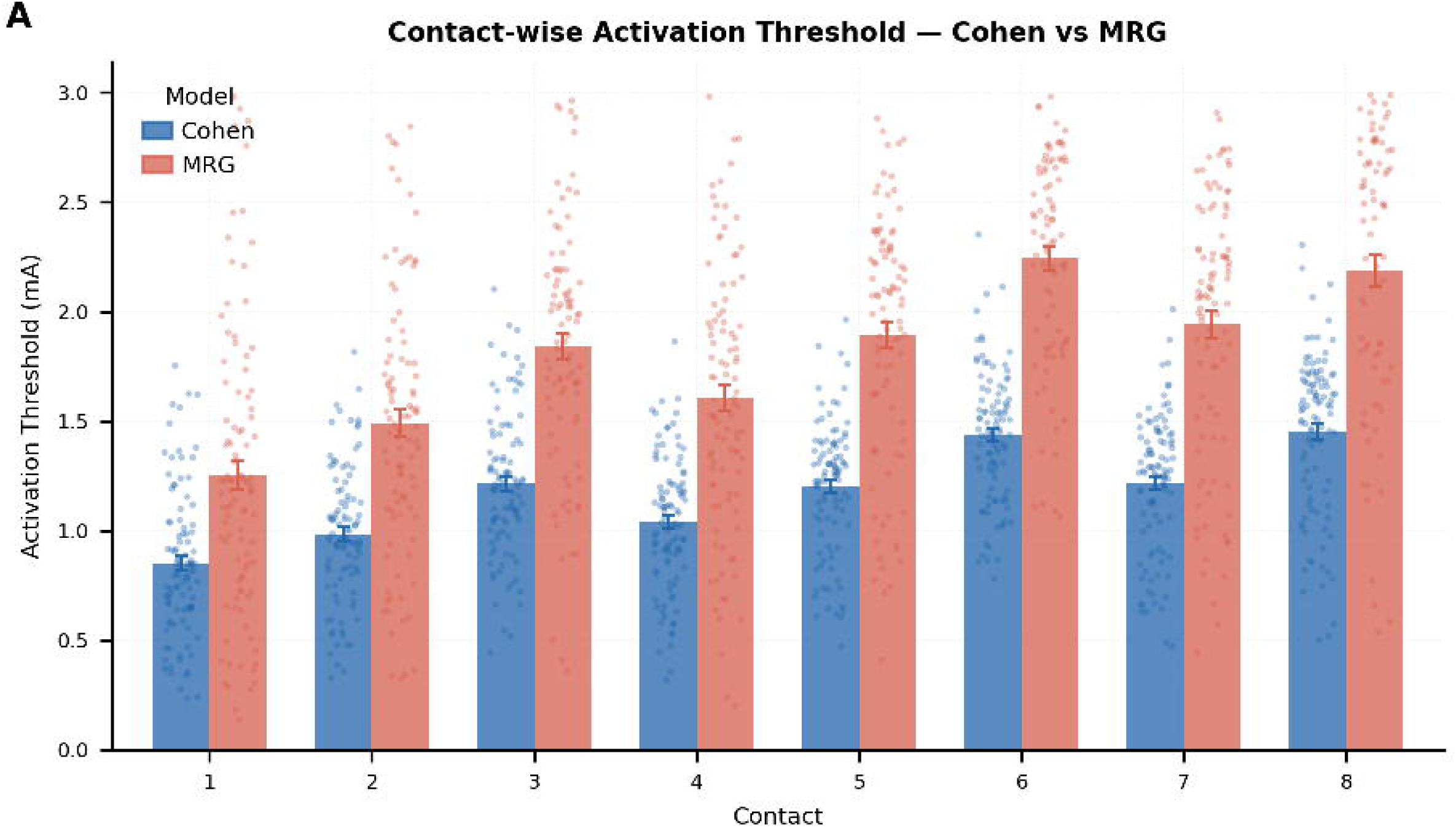
Comparison of mean activation thresholds across electrode contacts for Cohen and MRG models. Bar plot showing the mean activation threshold for each contact (1–8) predicted by the Cohen (blue) and MRG (red) models. Across all contacts, the MRG model consistently yields higher thresholds than the Cohen model. Both models exhibit similar spatial trends, with thresholds generally increasing from lower- to higher-numbered contacts, indicating systematic contact- dependent variation in activation sensitivity.

Figure 3 illustrates the relationship between activation threshold and the minimum distance to Contact 3 for both the Cohen and MRG models under biphasic pulse stimulation. In both models, activation threshold increased with increasing distance from the stimulating contact, demonstrating a strong distance-dependent effect on neural activation. The MRG model consistently exhibited higher activation thresholds and a steeper threshold–distance relationship than the Cohen model, indicating systematic differences in predicted excitability between the two formulations.

**Figure 3.**
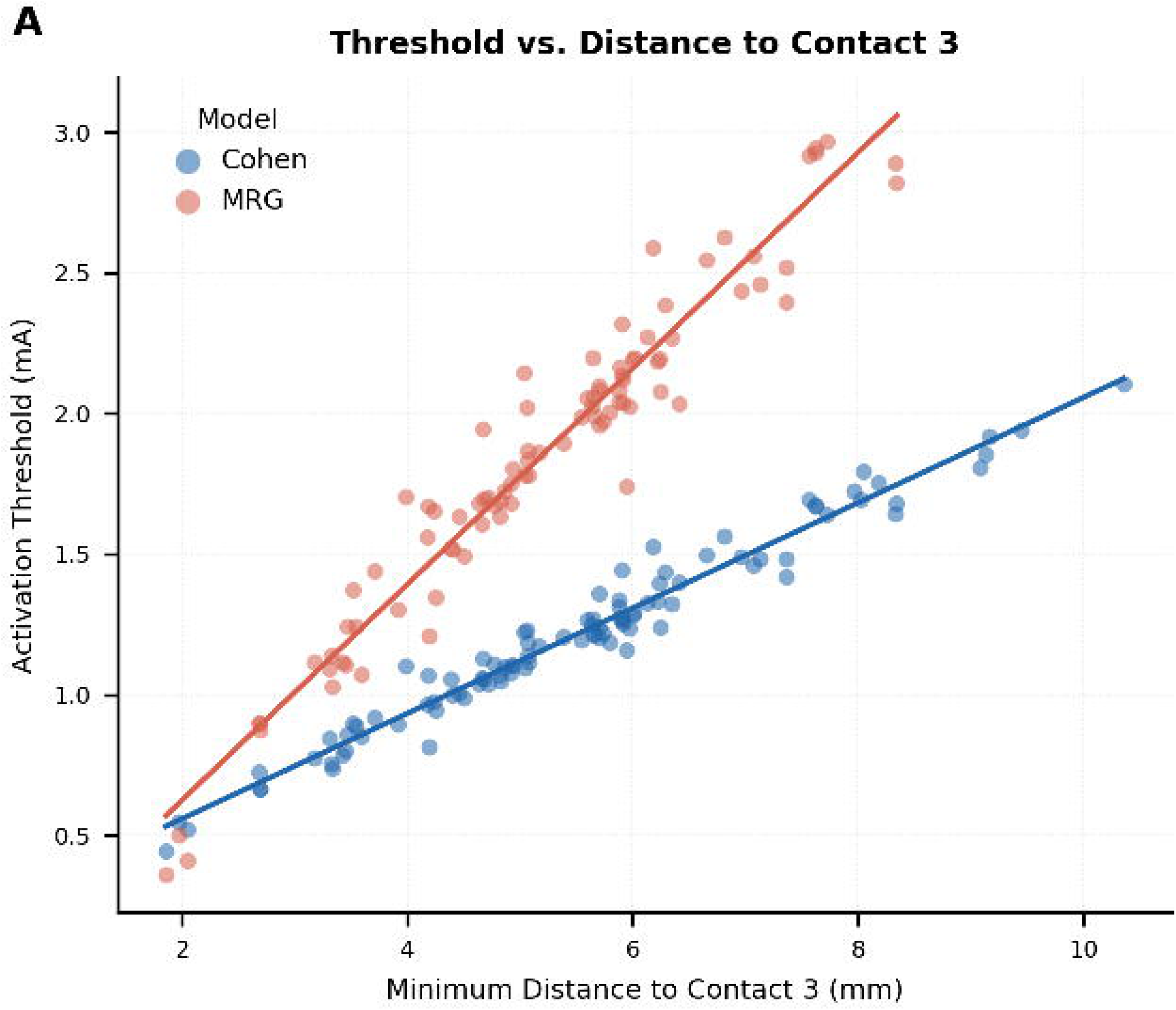
Threshold as a function of minimum distance to Contact 3 for Cohen and MRG models. Scatter plots showing activation threshold versus minimum fiber distance to Contact 3 for the two models (displayed in different colors). In both models, threshold increases monotonically with increasing distance, indicating a strong distance-dependent effect on activation. The MRG model exhibits consistently higher thresholds and a steeper distance–threshold relationship compared to the Cohen model, reflecting systematic differences in excitability predictions between formulations.

Figures 4 summarizes the relationship between activation threshold and stimulation parameters for model I and II. Both models exhibited a strong inverse relationship between pulse width and activation threshold: longer pulses reduced the required stimulation amplitude for activation. For the Cohen model, activation threshold decreased from approximately 1.56 at 60 μs to 1.18 at 120 μs, while the MRG model showed a similar trend, dropping from about 2.12 to 1.70 over the same range.

**Figure 4.**
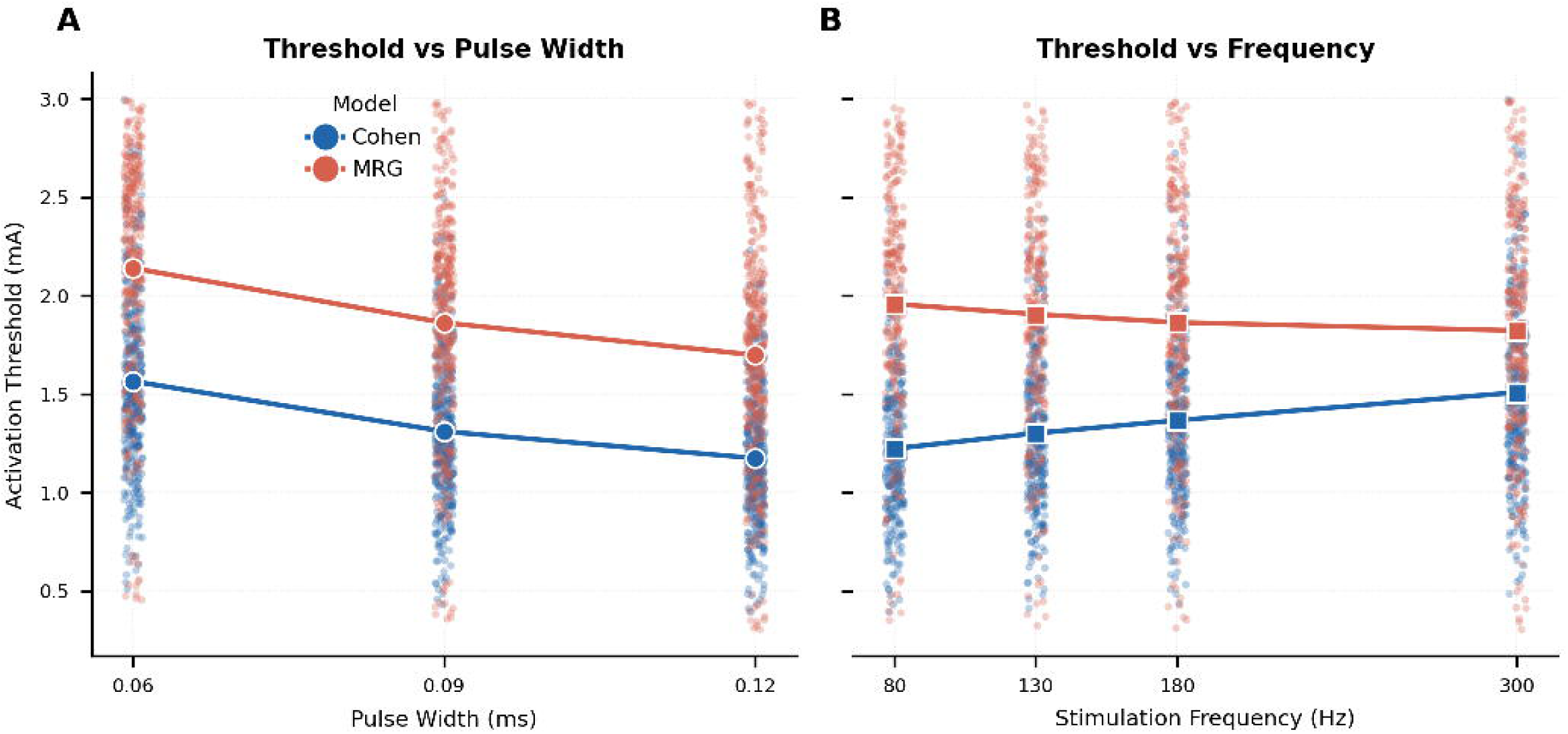
Mean activation threshold dependence on stimulation frequency and pulse width for MRG and Cohen models. >**(A)** Activation threshold vs. pulse width: Mean thresholds decrease with increasing pulse width for both models, with the MRG model consistently showing higher thresholds than the Cohen model. **(B)** Activation threshold vs. stimulation frequency: The MRG model shows a slight decrease in threshold with increasing frequency, whereas the Cohen model exhibits an increasing threshold trend with frequency.

However, the models differed in their response to stimulation frequency. The Cohen model demonstrated a positive correlation, with activation threshold increasing from roughly 1.23 at 80 Hz to 1.51 at 300 Hz. In contrast, the MRG model exhibited the opposite trend: activation threshold decreased from approximately 1.96 at 80 Hz to 1.82 at 300 Hz.

We compared the predictive performance of machine learning models for activation threshold estimation using model I and II. Gradient Boosting regression yielded the best results for both datasets. Model II achieved an R² of 0.986 on the test set, with a root mean squared error (RMSE) of 0.045 mA and a mean absolute error (MAE) of 0.034 mA (Table 1). In contrast, model I achieved an R² of 0.977, RMSE of 0.089 mA, and MAE of 0.068 mA. Cross-validation confirmed the robustness of both models, with CV R² values of 0.984±0.003 for model II and 0.975±0.004 for model I. These results indicate that both models can reliably predict activation thresholds based on distance, pulse width, and frequency, with the Cohen model providing slightly superior predictive performance.

**Table 1.** Performance metrics of linear regression, random forest, and gradient boosting models for predicting activation thresholds using the Cohen and MRG axon models. Metrics include the coefficient of determination (R²) for training and test sets, root mean squared error (RMSE), mean absolute error (MAE), and cross-validation mean and standard deviation.

| model | axon model | r2_train | r2_test | rmse_train | rmse_test | mae_test | cv_mean | cv_std |
| --- | --- | --- | --- | --- | --- | --- | --- | --- |
| Linear Regression | cohen | 0.9527 | 0.9435 | 0.0919 | 0.0902 | 0.0714 | 0.9512 | 0.0044 |
| Random Forest | cohen | 0.9960 | 0.9712 | 0.0268 | 0.0644 | 0.0475 | 0.9702 | 0.0046 |
| Gradient Boosting | cohen | 0.9901 | 0.9861 | 0.0420 | 0.0448 | 0.0337 | 0.9840 | 0.0025 |
| Linear Regression | mrg | 0.9281 | 0.9226 | 0.1567 | 0.1648 | 0.1230 | 0.9261 | 0.0094 |
| Random Forest | mrg | 0.9963 | 0.9767 | 0.0357 | 0.0905 | 0.0715 | 0.9686 | 0.0057 |
| Gradient Boosting | mrg | 0.9839 | 0.9772 | 0.0741 | 0.0895 | 0.0678 | 0.9750 | 0.0040 |

Figures 5 illustrates the relationships between activation threshold and key stimulation parameters for two models, respectively. In both models, the activation threshold increases strongly with distance from the electrode, confirming that spatial proximity is the primary determinant of neural activation. Pulse width and frequency exert secondary effects, with higher pulse widths and frequencies generally associated with slightly higher thresholds.

**Figure 5.**
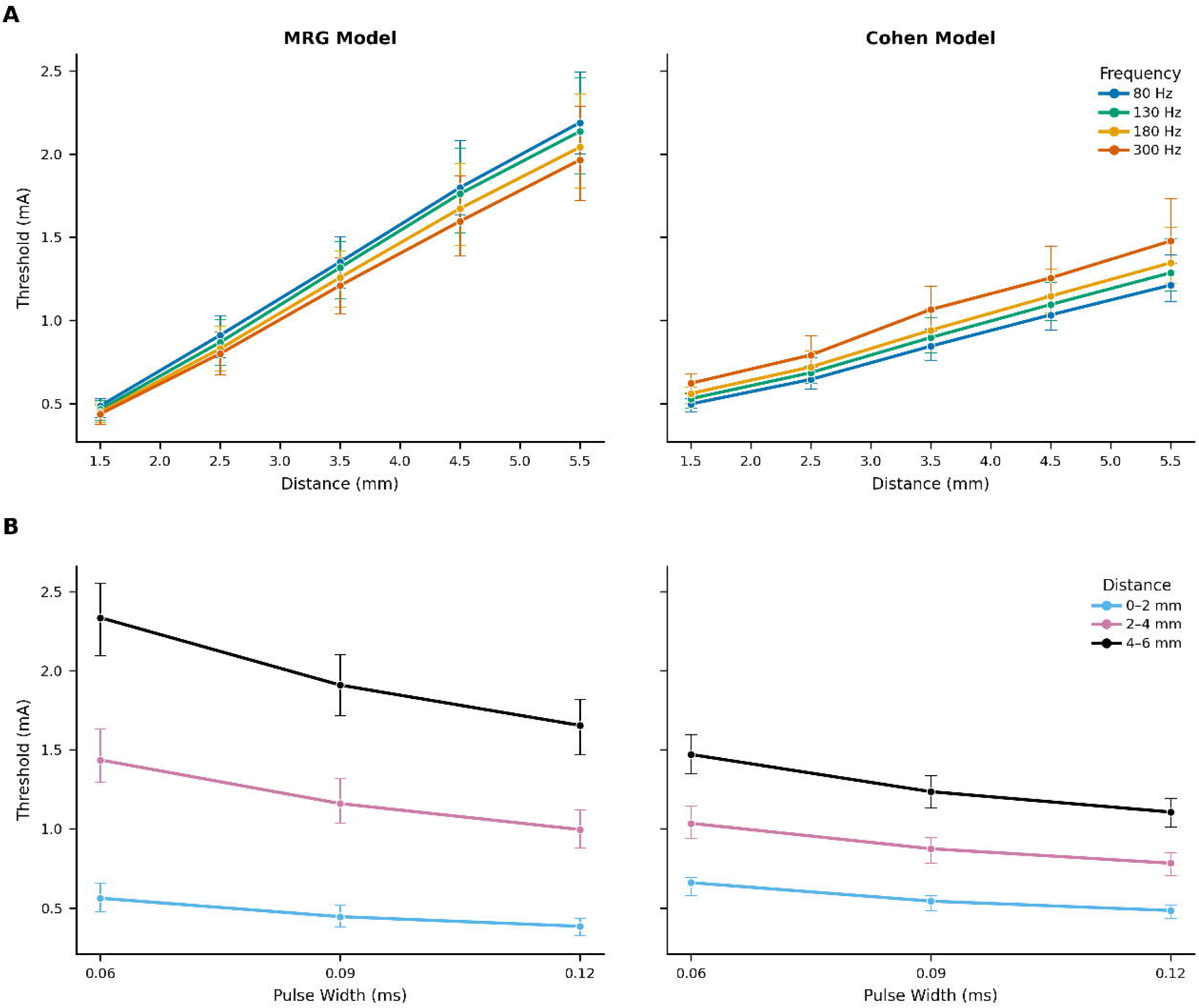
Effects of Distance, Pulse Width, and Frequency on Activation Thresholds in the Cohen and MRG Models. Model I (top row): (a) Threshold (mA) as a function of distance (mm) at 80, 130, 180, and 300 Hz, showing increasing threshold with distance across all frequencies. (b) Threshold (mA) versus pulse width (ms) for distance groups (0–2 mm, 2–4 mm, 4–6 mm), illustrating decreasing threshold with increasing pulse width and higher thresholds at greater distances. Model II (bottom row): (a) Threshold (mA) as a function of frequency (Hz) for the same distance groups, demonstrating modest frequency-dependent variations with consistently higher thresholds at larger distances. (b) Threshold (mA) versus distance (mm), colored by frequency, highlighting the dominant effect of distance on threshold and the secondary modulation by stimulation frequency.

Both models demonstrated strong agreement in predicting activation thresholds (R² = 0.780, Pearson r = 0.883, p < 0.001) (Figure 6). The linear regression equation was:

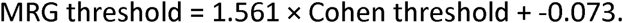

**Figure 6.**
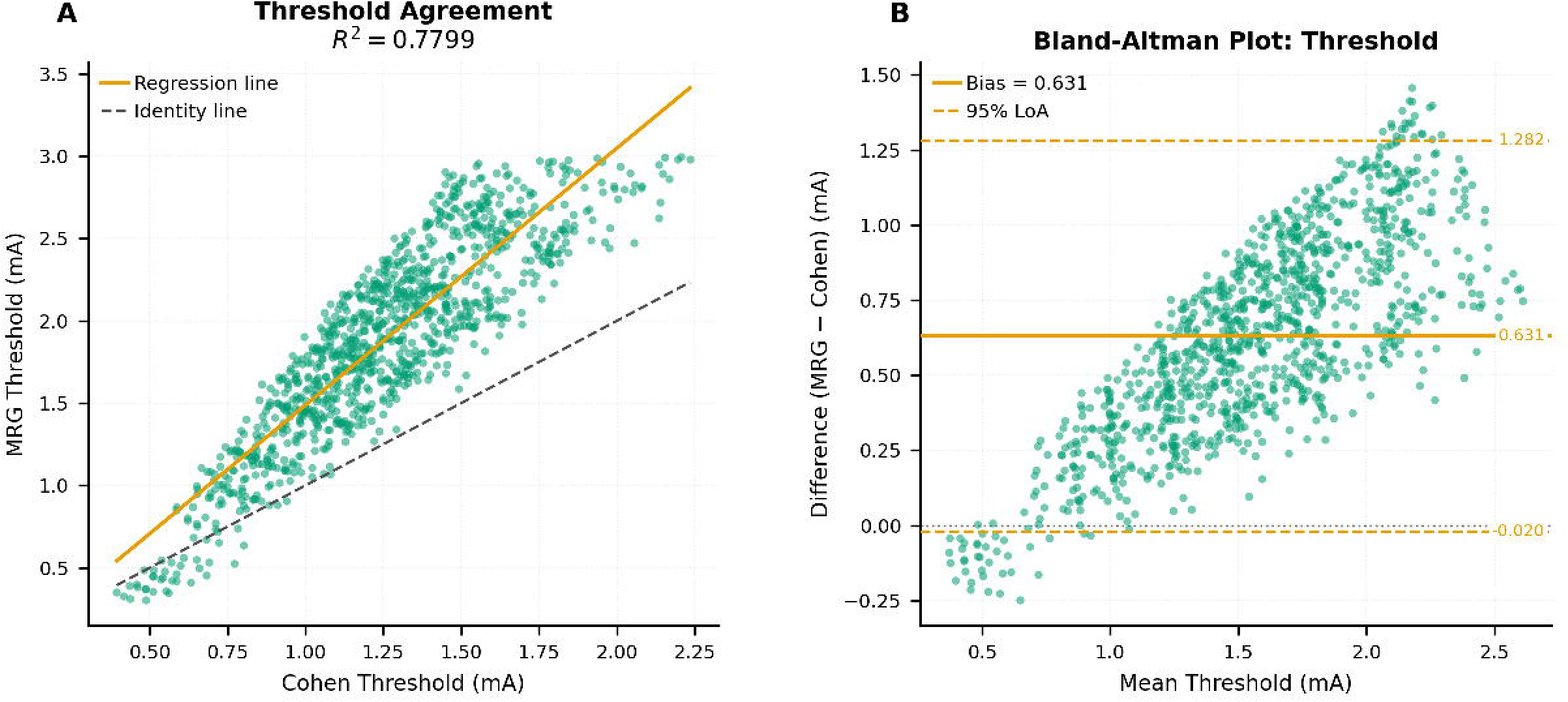
Agreement analysis between Cohen and MRG threshold estimates. (a) Scatter plot of MRG versus Cohen thresholds (mA) with linear regression (red) and identity (dashed) lines, demonstrating strong correlation (R² = 0.7799, Pearson r = 0.8831) and a systematic deviation from unity. (b) Bland–Altman plot showing the difference (MRG − Cohen) versus mean threshold, indicating a positive bias of 0.6311 mA and 95% limits of agreement from −0.0201 to 1.2823 mA.

The RMSE was 0.275 mA and MAE was 0.230 mA. Bland-Altman analysis revealed a mean bias of 0.631 mA (95% limits of agreement: −0.020 to 1.282 mA), indicating that the MRG model predicted higher thresholds on average compared to the Cohen model.

## 4. Discussion

In this study, we compared two neuron models for their applicability in DBS simulations. These models were selected because they represent different biophysical approaches to modeling neuronal responses, allowing us to evaluate how these differences influence predicted DBS activation thresholds and sensitivity to stimulation parameters.

Both models are capable of simulating repetitive stimulation pulses. This enables representation of refractory periods and cumulative temporal effects, providing a closer approximation to real-world DBS conditions in which stimulation is delivered continuously rather than as isolated pulses.

Our results show that both models were sensitive to variations in pulse width and stimulation frequency, confirming that these parameters play a critical role in activation dynamics. A systematic difference in activation thresholds was observed, with Cohen model generally requiring lower thresholds than MRG model, which highlights biophysical differences between the two modeling approaches.

Importantly, within both models, axon–electrode distance had a stronger influence on activation threshold than pulse width or frequency. Although pulse width and frequency affected activation thresholds, their impact was smaller than that of axon–electrode distance. This result, supported by the machine learning analysis, highlights the importance of accurate spatial information for reliable threshold prediction.

We also observed opposite effects of increasing stimulation frequency across two models. In Cohen model, the activation threshold increased with higher frequencies, likely due to the presence of a refractory period. Milosevic et al. [41, 42] investigated the influence of stimulation frequency and synaptic area across different brain regions and reported that neuronal activity within the STN progressively decreases as stimulation frequency rises. Although synaptic effects were not explicitly modeled in our work, we observed a similar trend of reduced activation at higher frequencies in Cohen model, which contrasts with the predictions of MRG model.

These differences in activation behavior can be traced back to the distinct structural and electrical representations of the axon in the two models. One key difference between the models is their double-cable structure. Cohen model considers a nanocircuit under the myelin, whereas MRG model treats the internode as much more electrically isolated. In MRG model, the myelin sheath is represented using a capacitance and small leak conductance and the periaxonal space is assumed to be tightly sealed and grounded at the node. This means internodal regions cannot sustain significant longitudinal current, so most charging and discharging happens through nodal transmembrane pathways. In contrast, Cohen model uses biophysically realistic myelin built from actual wrap count (15 layers), includes finite myelin conductance, and does not assume perfect paranodal sealing. Instead, a thin but conductive periaxonal layer (∼50 Ω·cm) forms a continuous resistive channel beneath the sheath, with paranodes modeled as high-resistance but incomplete seals. This creates a sub-myelin nanocircuit that allows depolarizing current to propagate along the internode and feed back into neighboring nodes. Consequently, activation spreads via dual pathways (axoplasm and periaxonal space) resulting in greater excitability, lower activation thresholds, and a more realistic node-internode voltage differential during action potential propagation [20, 43].

Another key difference between the models is ion channel implementation. Both models are built on physiological data but implemented in slightly different ways. At the nodes of Ranvier, MRG model incorporates fast sodium (Na⁺) channels for action potential initiation and propagation, delayed rectifier potassium (K⁺) channels for repolarization, and persistent sodium and slow potassium channels that influence afterpotentials and recovery cycle dynamics. The paranodal (MYSA, FLUT) and internodal (STIN) segments use passive (pas) mechanisms with leak conductances but no active voltage-gated channels, reflecting the electrical insulation provided by myelin. Cohen Model incorporates a more detailed representation of ion channel dynamics compared to MRG model. It includes multiple sodium and potassium channel subtypes, such as fast sodium currents (Na, Na_is_, Na_x_) and a persistent sodium current (Na_p_), as well as diverse potassium conductances: a delayed rectifier (K_v_), low-threshold K_v1_ channels, a specialized axonal K_v1_ channel (K_v1,ax_), a K_v7_/M-type channel (K_v7_), and a calcium-activated potassium channel (K_ca_). This modular architecture enables the model to capture nuanced mechanisms of action potential initiation, propagation, and adaptation, offering a more biophysically realistic framework for DBS simulations.

Finally, an important structural distinction between the models lies in the representation of axonal diameter. Cohen model incorporates variable diameters across nodes, which may provide a more physiologically realistic representation of axonal heterogeneity [43–45]). In contrast, MRG model assumes a constant diameter across nodes, simplifying the geometry but potentially reducing biological fidelity. Additionally, MRG model relies on older physiological data, which may reduce accuracy under current standards. Updating the model with more recent measurements of axonal properties could improve its physiological relevance and predictive performance [46].

In addition to structural and biophysical differences, computational complexity is another important consideration. Cohen model tends to be more computationally demanding due to its inclusion of detailed parameters for myelin and submyelin spaces, whereas MRG model is simpler but still highly effective for many DBS simulations. This trade-off between realism and efficiency should guide model selection based on the goals and resources of a given study.

Several limitations of this study should be noted. First, Cohen model incorporates soma and dendritic compartments in addition to the axon, whereas MRG model is restricted to axonal segments. While both models are appropriate for studying stimulation-induced axonal activation, differences in morphological complexity could influence quantitative predictions.

Differences in morphological complexity between the models may contribute to variations in predicted excitability and frequency-dependent behavior, although axonal activation remains the primary determinant of DBS thresholds.

Second, errors introduced by coregistration, localization, and the relatively low spatial resolution of diffusion-weighted imaging (DWI) data, along with the assumption of identical tissue conductivity values for both models, may affect the absolute accuracy of fiber tracking, model alignment, and current spread estimation. Future work should or could? employ higher-resolution acquisitions, advanced reconstruction techniques, and region-specific conductivity estimates to improve physiological realism. Nevertheless, because both models were evaluated under identical imaging, registration, and conductivity conditions, these limitations are unlikely to bias the comparative conclusions drawn in this study.

Third, this study was conducted on a single patient to highlight model differences and avoid the added complexity of multi-patient variability. Future studies should evaluate these models across larger patient cohorts to ensure generalizability. Additionally, incorporating indirect measurements such as EMG recordings could help correlate model predictions with clinical outcomes. Ultimately, in vivo validation remains essential for confirming model predictions [47].

## 5. Conclusion

This study systematically compared the McIntyre-Richardson-Grill model and Cohen model in deep brain stimulation simulations. Both models were sensitive to key stimulation parameters and achieved high predictive accuracy when combined with machine learning approaches. However, consistent differences were observed: the Cohen model exhibited lower activation thresholds, greater excitability across most tracts, and a frequency-dependent increase in threshold, whereas the MRG model predicted higher thresholds and showed the opposite frequency trend. These discrepancies likely reflect fundamental structural and biophysical differences, including myelin representation, periaxonal conductivity, and ion channel dynamics. Despite these differences, overall agreement between the models remained strong, supporting their reliability for estimating DBS-induced axonal activation. Electrode distance emerged as the primary determinant of activation threshold, with pulse width and frequency exerting secondary effects. Model selection should therefore balance computational efficiency and biophysical realism according to the specific requirements of the application, and further validation with larger patient cohorts and in vivo data will be essential to advance patient- specific DBS modeling.

## Footnotes

1 https://modeldb.science/3810

2 https://modeldb.science/260967

